# Superficial white matter microstructure affects processing speed in cerebral small vessel disease

**DOI:** 10.1101/2021.12.30.474604

**Authors:** Shuyue Wang, Fan Zhang, Peiyu Huang, Hui Hong, Yeerfan Jiaerken, Xinfeng Yu, Ruiting Zhang, Qingze Zeng, Yao Zhang, Ron Kikinis, Yogesh Rathi, Nikos Makris, Min Lou, Ofer Pasternak, Minming Zhang, Lauren J. O’Donnell

**Affiliations:** Department of Radiology, the Second Affiliated Hospital of Zhejiang University School of Medicine, China; Brigham and Women’s Hospital, Harvard Medical School, Boston, Massachusetts, USA; Center for Morphometric Analysis, Massachusetts General Hospital, Harvard Medical School, Boston, Massachusetts, USA; Department of Neurology, the Second Affiliated Hospital of Zhejiang University School of Medicine, China

**Keywords:** superficial white matter, white matter hyperintensities, extracellular free-water, processing speed, diffusion magnetic resonance imaging, cerebral small vessel disease

## Abstract

White matter hyperintensities (WMH) are a typical feature of cerebral small vessel disease (CSVD). This condition contributes to about 50% of dementias worldwide, a massive health burden in aging. Microstructural alterations in the deep white matter (DWM) have been widely examined in CSVD. However, little is known about abnormalities in the superficial white matter (SWM) and their relevance for processing speed, the main cognitive deficit in CSVD. In this paper, 141 patients with CSVD were studied. Processing speed was assessed by the completion time of the Trail Making Test Part A. White matter abnormalities were assessed by WMH burden (lesion volume on T2-FLAIR) and diffusion MRI, including DTI and free-water (FW) imaging microstructure measures. The results of our study indicate that the superficial white matter may play a particularly important role in cognitive decline in CSVD. SWM imaging measures resulted in a large contribution to processing speed, despite a relatively small WMH burden in the SWM. SWM FW had the strongest association with processing speed among all imaging markers and, unlike the other diffusion MRI measures, significantly increased between two patient subgroups with the lowest WMH burdens (possibly representing early stages of disease). When comparing two patient subgroups with the highest WMH burdens, the involvement of WMH in the SWM was accompanied by significant differences in processing speed and white matter microstructure. Given significant effects of WMH volume and regional FW on processing speed, we performed a mediation analysis. SWM FW was found to fully mediate the association between WMH volume and processing speed, while no mediation effect of DWM FW was observed. Overall, our findings identify SWM abnormalities in CSVD and suggest that the SWM has an important contribution to processing speed. Results indicate that FW in the SWM is a sensitive marker of microstructural changes associated with cognition in CSVD. This study extends the current understanding of CSVD-related dysfunction and suggests that the SWM, as an understudied region, can be a potential target for monitoring pathophysiological processes in future research.

## 1. Introduction

Cerebral small vessel disease (CSVD) is the most important vascular contributor to cognitive decline and the cause of nearly 50% of dementia worldwide (Wardlaw, Smith and Dichgans, 2019). CSVD-related cognitive impairment is often characterized as deficits in executive function and speed of information processing (Duering *et al*., 2014; Olivia K. L. Hamilton *et al*., 2021). White matter hyperintensities (WMH), the most prevalent feature of CSVD on magnetic resonance imaging (MRI) (Vergoossen *et al*., 2021), appear as hyper-intense patches in the white matter on T2-weighted (T2w) and T2-FLAIR images (Fazekas *et al*., 1987; Wardlaw *et al*., 2013). Widespread microstructural changes in the white matter are a key manifestation in CSVD and have been consistently associated with cognitive deficits (Cremers *et al*., 2016; Olivia K. L. Hamilton *et al*., 2021; Vergoossen *et al*., 2021).

The most commonly used method to study the white matter microstructure is diffusion MRI (dMRI) (Basser, Mattiello and LeBihan, 1994), which measures diffusion properties of water molecules in vivo and in a non-invasive way. dMRI studies have found widespread white matter microstructural alterations and significant associations with cognition in CSVD (Table 1). Many studies have focused on conventional diffusion measures, where a decrease in the anisotropy of water diffusion (fractional anisotropy, FA) and an increase in the extent of water diffusion (mean diffusivity, MD) are generally considered indicative of damaged white matter microstructure (van Norden *et al*., 2012; Baykara *et al*., 2016). Using these conventional diffusion measures, previous studies have suggested that potential pathological processes underlying WMH may be related to demyelination and reduced axonal density (Wardlaw, Valdés Hernández and Muñoz-Maniega, 2015; Muñoz Maniega *et al*., 2017). Furthermore, recent studies indicate that an increase of extracellular free-water (FW) (Pasternak *et al*., 2009), measured using dMRI, may be an early pathological process in CSVD (Duering *et al*., 2018; Wardlaw, Smith and Dichgans, 2019; Humphreys, Smith and Wardlaw, 2021) and is a sensitive marker of cognitive performance in aging (Maillard *et al*., 2019). While associations between white matter microstructure and cognitive performance are found throughout the white matter in CSVD (Table 1), thus far the superficial white matter microstructure has not been specifically investigated.

**Table 1:** Summary of dMRI studies that explore associations between white matter microstructure and cognitive performance (processing speed or executive function) in cerebral small vessel disease. For each study, the methodology used to evaluate white matter microstructure and the locations of findings are reported.

| CSVD dMRI study methodology |  |
| --- | --- |
| Conventional diffusion measures (e.g., FA, MD) | (Michael O’Sullivan *et al.*, 2004; M. O’Sullivan *et al.*, 2004; O’Sullivan *et al.*, 2005; Nitkunan *et al.*, 2008; Schmidt *et al.*, 2010; Quinque *et al.*, 2012; Lawrence *et al.*, 2013; Tuladhar *et al.*, 2015; Baykara *et al.*, 2016; Moonen *et al.*, 2017; Zeestraten *et al.*, 2017; Tozer *et al.*, 2018; Chen *et al.*, 2019; Wei *et al.*, 2019; Huang *et al.*, 2020; Liu *et al.*, 2020; Du *et al.*, 2021; Vergoossen *et al.*, 2021) |
| Advanced diffusion measures (e.g. FW, NODDI, DKI, IVIM) | (Zhang, Wong, van de Haar, *et al.*, 2017; Duering *et al.*, 2018; Gesierich *et al.*, 2020; Konieczny *et al.*, 2021) |
| Structural network measures | (Lawrence *et al.*, 2014; Reijmer *et al.*, 2016; Tuladhar *et al.*, 2016, 2017; Heinen *et al.*, 2018; van Leijsen *et al.*, 2019; Boot *et al.*, 2020; Gesierich *et al.*, 2020; Liu *et al.*, 2020; Du *et al.*, 2021) |
| **Study findings: White matter regions related to cognitive performance in CSVD** |  |
| **Global findings** |  |
| In the entire white matter or entire white matter skeleton | (Nitkunan *et al.*, 2008; Baykara *et al.*, 2016; Moonen *et al.*, 2017; Zeestraten *et al.*, 2017; Wei *et al.*, 2019; Du *et al.*, 2021; Konieczny *et al.*, 2021) |
| **Localized findings** |  |
| Within WMH | (Michael O’Sullivan *et al.*, 2004; M. O’Sullivan *et al.*, 2004; Schmidt *et al.*, 2010) |
| In normal-appearing white matter | (Michael O’Sullivan *et al.*, 2004; M. O’Sullivan *et al.*, 2004; Schmidt *et al.*, 2010; Jokinen *et al.*, 2013; Lawrence *et al.*, 2013, 2014; Tuladhar *et al.*, 2015; Zhang, Wong, Uiterwijk, *et al.*, 2017; Tozer *et al.*, 2018) |
| In particular fiber tracts or deep white matter regions | (O’Sullivan *et al.*, 2005; Quinque *et al.*, 2012; Tuladhar *et al.*, 2015; Chen *et al.*, 2019; Huang *et al.*, 2020; Liu *et al.*, 2020; Vergoossen *et al.*, 2021) |
| **Network findings** |  |
| Global network | (Lawrence *et al.*, 2014; Tuladhar *et al.*, 2016; Heinen *et al.*, 2018; van Leijsen *et al.*, 2019; Gesierich *et al.*, 2020; Du *et al.*, 2021) |
| Local (sub)network | (Lawrence *et al.*, 2014; Reijmer *et al.*, 2016; Boot *et al.*, 2020; Liu *et al.*, 2020; Du *et al.*, 2021) |
| Peripheral white matter connections | (van Leijsen *et al.*, 2019) |

**Abbreviations:** CSVD, cerebral small vessel disease; TBSS, tract-based spatial statistics; FA, fractional anisotropy; MD, mean diffusivity; FW, free water; NODDI, neurite orientation dispersion and density imaging; DKI, diffusion kurtosis imaging, IVIM, intravoxel incoherent motion imaging; WMH, white matter hyperintensity.

The superficial white matter (SWM) is the thin layer of WM underneath the cortex. It includes short-range association connections (u-fibers) that connect adjacent gyri, and this region is also traversed by terminations of long-range tracts (Liu *et al*., 2016). Autopsy and MRI studies have reported that the short association fibers make up 57-67% of all white matter fibers, and their estimated volume is much larger than the volume of the long-range projections (Schüz and Braitenberg, 2002; Vergani *et al*., 2014; Liu *et al*., 2016). The SWM plays an important role in processing speed and other functions such as working memory and visuomotor attention (Nazeri *et al*., 2013, 2015; Phillips *et al*., 2018). In recent years, dMRI studies have suggested the importance of the SWM in aging (Phillips *et al*., 2013; Nazeri *et al*., 2015; Cox *et al*., 2016), Alzheimer’s disease (Phillips *et al*., 2016; Bigham *et al*., 2020; Contarino *et al*., 2021), and neuropsychiatric disease (D’Albis *et al*., 2017; d’Albis *et al*., 2018). In CSVD, dMRI tractography studies of the entire brain as a network have implicated the SWM (Table 1): two studies found reduced SWM connectivity (in short peripheral tracts (Tuladhar *et al*., 2017) and some superficial white matter u-fibers (Lawrence *et al*., 2014)), while one study found that a decline in peripheral connection strength was associated with cognition (van Leijsen *et al*., 2019). However, despite the well-known decline in processing speed (Prins *et al*., 2005; Peng, Geriatric Neurology Group, Chinese Society of Geriatrics and Clinical Practice Guideline for Cognitive Impairment of Cerebral Small Vessel Disease Writing Group, 2019; O. K. L. Hamilton *et al*., 2021), which is associated with widespread white matter changes in CSVD (Konieczny *et al*., 2021; Vergoossen *et al*., 2021), the abnormalities in the SWM are poorly understood.

Therefore, we believe that studying white matter abnormalities in the SWM may shed light on the mechanisms underlying cognitive dysfunction in CSVD. Several lines of evidence suggest that the SWM may be affected. First, the effect of WMH is widespread. A recent review summarizes postmortem histological and MR imaging studies of CSVD, which have reported extensive white matter lesions with SWM involvement (Humphreys, Smith and Wardlaw, 2021). In addition, progression of WMH is known to spread from deeper to more superficial white matter regions during the course of CSVD (Lambert *et al*., 2016; van Leijsen *et al*., 2018).

In this study, our primary goal is to employ imaging markers to investigate the impact of the SWM on processing speed, the main cognitive deficit in CSVD (O. K. L. Hamilton *et al*., 2021; Vergoossen *et al*., 2021). First, we assess the effect of WMH burden on the SWM and DWM microstructure. Second, we assess the contribution of abnormalities in the SWM and the DWM towards processing speed. Third, we assess differences of SWM and DWM microstructure and processing speed between groups with different levels of WMH burden. Fourth, we perform mediation analysis to further determine whether SWM and DWM microstructure could mediate the association between WMH burden and processing speed.

## 2. Materials and methods

### 2.1 Subjects

This study was approved by the Medical Ethics Committee of the Second Affiliated Hospital, Zhejiang University School of Medicine. Written informed consent was obtained from each subject.

We recruited patients admitted to the Department of Neurology, the Second Affiliated Hospital, Zhejiang University School of Medicine, who received brain MRI and were diagnosed with CSVD between December 2015 and December 2020. More details about the cohort can be found in previously published studies (Wang *et al*., 2020; Huang, Zhang, Jiaerken, Wang, Hong, *et al*., 2021). Visual assessment of WMH presence (Fazekas scores) was performed by 2 neuroradiologists (R.Z. and H.H) according to the STandards for ReportIng Vascular changes on nEuroimaging (STRIVE) (Wardlaw *et al*., 2013). Inclusion criteria were as follows: (a) visible WMH on T2-FLAIR; (b) age > 40; (c) normal vision and hearing. Exclusion criteria were as follows: (a) WM lesions of nonvascular origin (immunological-demyelinating, metabolic, toxic, infectious, etc.); (b) severe head motion during MRI scanning; (c) history of stroke, multiple sclerosis, Alzheimer’s disease, Parkinson’s disease, or head trauma; (d) image quality issues. Starting from a total of 365 subjects who had MRI data, we first excluded 63 subjects with possible other brain disorders under the exclusion criteria. Of the remaining 302, there were 145 subjects with the processing speed assessment. 4 subjects were further excluded due to insufficient image quality. Finally, a total of 141 subjects were included in the present study. Demographic information and vascular risk factors, including age, sex, diabetes, hypertension, hyperlipidemia, heart disease, smoking, and drinking histories, are summarized in Table 1.

### 2.2 Neuropsychological assessments

Each subject’s neuropsychological condition was assessed by the mini-mental state examination (MMSE) and Montreal Cognitive Assessment (MoCA). Processing speed was evaluated with the Trail Making Test Part A (TMT-A) (Bowie and Harvey, 2006). The completion time for TMT-A was evaluated by scoring the time in seconds, and the maximum was limited to 180s according to the administration procedure and interpretive guideline (Thompson *et al*., 1999; Bowie and Harvey, 2006). Then, the TMT-A completion time was log-transformed to reduce the skewness (Yano *et al*., 2017; Sudre *et al*., 2019).

### 2.3 MRI data acquisition and preprocessing

All subjects underwent multi-modal MRI on a 3.0 T MR scanner (MR750, GE Healthcare, Milwaukee, USA) equipped with an 8-channel brain phased array coil. The scanned modalities for each subject included T1-weighted imaging (T1w), T2-FLAIR imaging and dMRI. T1-weighted imaging was acquired using spoiled gradient echo sequences with TR/TE = 7.3/3.0 ms, TI = 450 ms, flip angle = 8°, slice thickness = 1 mm, matrix = 250 × 250, field of view (FOV) = 250 mm × 250 mm. The sequence parameters of T2-FLAIR were: TR/TE = 8,400/152 ms, TI = 2,100 ms, flip angle = 90°, slice thickness = 4 mm without slice gap, matrix size = 256 × 256, FOV = 240 mm × 240 mm. dMRI was acquired with a single shot, diffusion-weighted spin echo echo-planar imaging sequence. 30 noncollinear directions were acquired with a b-value of 1,000 s/mm^2^. 5 additional volumes were acquired without diffusion weighting (b-value = 0 s/mm^2^). Other parameters of dMRI were as follows: TR/TE = 8,000/80.8 ms, flip angle = 90°, slice thickness = 2 mm without slice gap, isotropic voxel size = 2 × 2 × 2 mm^3^, matrix size = 128 × 128, FOV = 256 mm × 256 mm. All dMRI data was preprocessed using MRtrix3 (http://www.mrtrix.org) (Tournier *et al*., 2019), including signal denoising, Gibbs ringing removal, eddy-current and head motion correction, and bias field correction.

### 2.4 Image processing and estimation of imaging markers

Figure 1 gives an overview of the image processing steps for the computation of imaging markers of interest. We computed two types of imaging markers in the SWM, including 1) WMH volume to assess the burden of WMH and 2) diffusion measures to assess microstructural abnormalities. For comparison, we also computed the imaging markers in the DWM. The major image processing steps included (a) regional white matter segmentation, (b) WMH segmentation, and (c) diffusion measure computation, as described below in detail.

**Figure 1.**
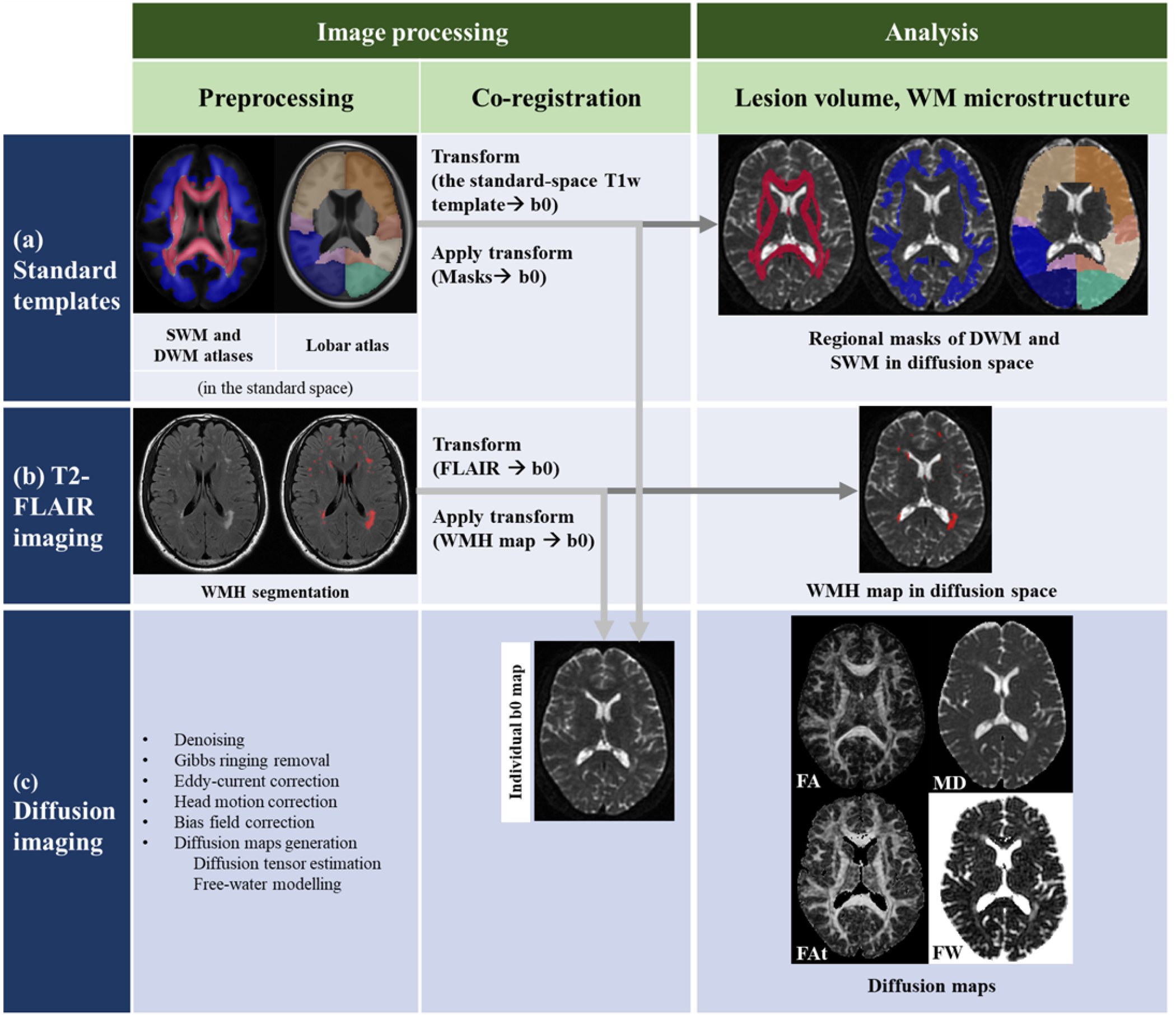
Schematic illustration of the multi-modal image processing and analysis pipeline. *Abbreviations:* SWM, superficial white matter; DWM, deep white matter; WMH, white matter hyperintensity; CSF, cerebrospinal fluid.

#### 2.4.1 White matter segmentation

The entire white matter was segmented into deep white matter (DWM) and superficial white matter (SWM). This was done by registering DWM and SWM atlases defined in the standard MNI space to each subject’s diffusion MRI space. Specifically, DWM was defined as the white matter region comprising the JHU-ICBM-DTI-81 white matter atlas (Mori *et al*., 2005). SWM was defined using an atlas of the SWM across the adult lifespan (N=141; 18-86 years of age) (Nazeri *et al*., 2015). This SWM atlas was created to be used together with the JHU DWM atlas; thus, it excludes the DWM regions. It has been successfully applied to study the relationship between diffusion measures in SWM and cognitive performance in aging (Nazeri *et al*., 2015). The standard-space DWM and SWM masks were aligned to each subject’s diffusion space via a nonlinear registration between the standard-space T1w template and the subject’s b0 image using the Normalization toolbox in Statistical Parametric Mapping (SPM12, http://www.fil.ion.ucl.ac.uk/spm). All registered masks were visually inspected in each subject. We also further segmented the SWM according to the brain lobes to investigate lobar SWM abnormalities. Brain lobes were defined using a standard atlas (Mayo Clinic Adult Lifespan Template, MCALT, www.nitrc.org/projects/mcalt) (Schwarz *et al*., 2018). A total of 8 lobar regions were studied, including the bilateral frontal, occipital, temporal, and parietal lobes. The standard-space lobar masks were aligned to each subject’s diffusion space via a registration between the standard-space T1w template and the subject’s b0 image. Image registration was conducted using SPM. After visual inspection for registered lobar masks, lobar SWM regions were defined according to the subject’s lobar masks and the SWM mask.

#### 2.4.2 WMH segmentation and WMH volume computation

White matter hyper-intense (WMH) regions were segmented using the T2-FLAIR images. The lesion prediction algorithm (LPA) of the Lesion Segmentation Toolbox (http://www.applied-statistics.de/lst.html) in SPM was used to automatically segment WMH lesions in the T2-FLAIR image of each subject under study. Previous studies (Cedres *et al*., 2020; Vanderbecq *et al*., 2020; Wang *et al*., 2020; Zhang *et al*., 2021) have shown successful applications of the LPA algorithm in WMH segmentation using T2-FLAIR. Each segmented result was visually inspected and manually corrected by expert readers (S.W. and H.H.) in ITK-SNAP (http://www.itksnap.org) according to the STandards for ReportIng Vascular changes on nEuroimaging (STRIVE) (Wardlaw *et al*., 2013). One neuroradiologist (R.Z.) reviewed all the corrected lesion maps. Then, the segmented WMH lesion map was registered to the individual diffusion space by performing a registration between the T2-FLAIR image and the first b0 diffusion image using SPM.

To assess the burden of disease, we measured the volume of WMH (the larger the volume, the heavier the burden of disease). The volume of the segmented WMH region (referred to as the WMH-total volume), as well as the volume of WMH in the SWM and DWM regions (referred to as the WMH-SWM and WMH-DWM volumes, respectively), were calculated for each subject using the *fslstats* tool in the FMRIB Software Library (FSL, https://fsl.fmrib.ox.ac.uk/fsl/fslwiki).

#### 2.4.3 Diffusion measure computation

Four diffusion measures of interest were computed from the DWI data, including: conventional fractional anisotropy (FA) and mean diffusivity (MD) (Pierpaoli and Basser, 1996), and extracellular free-water (FW) and FW-corrected intracellular tissue FA (FAt) (Pasternak *et al*., 2009). FA and MD were computed using the conventional diffusion tensor model (Pierpaoli and Basser, 1996). This was done by fitting the preprocessed DWI data to a diffusion tensor image using a linear regression in the FSL *dtifit* tool. FA and MD images were then computed from the estimated diffusion tensor in each voxel using the FSL *fslmaths* tool. FW and FAt were computed using free-water (FW) imaging (Pasternak *et al*., 2009). FW imaging is a model-based approach that applies a regularization framework to fit a two-compartment model to diffusion-weighted images. The two compartments estimated in the FW model are: 1) an isotropic FW compartment, which quantifies the contribution of extracellular FW to the signal, and 2) a tissue compartment, modeled using a single diffusion tensor, which is corrected for FW. FW provides a putative index of unrestricted extracellular water content. FAt is calculated from the FW-corrected intracellular tissue compartment and more closely reflects changes in myelination and axonal membrane health than the conventional DTI metric FA (Pasternak *et al*., 2009, 2018). FW modeling was performed using a non-linear regularized minimization process implemented in MATLAB as previously described (Pasternak *et al*., 2009; Di Biase *et al*., 2020). The fractional volume of the FW compartment was estimated to produce a FW map. Then FAt map was calculated from the FW-corrected tensor using FSL.

The mean FA, MD, FW and FAt in the DWM and SWM were calculated for each subject using the FSL *fslstats* tool. In the same way, the mean of each diffusion measure was calculated for each lobar SWM region.

#### 2.4.4 Estimation of intracranial volume

T1-weighted images were processed using the default “Recon-all” pipeline in the FreeSurfer software (version 6.0, http://surfer.nmr.mgh.harvard.edu/fswiki). The intracranial volume (ICV) of each subject was calculated and extracted as a covariate.

### 2.5 Statistical analyses

We performed four analyses to investigate abnormalities in SWM and their effects on processing speed in the CSVD patients under study. We also performed the corresponding analyses in the DWM for comparison. All statistical analyses were performed in SPSS (version 23.0), R (version 4.0.5) and MATLAB (version R2019b). Covariates included age, sex, and ICV, with the addition of years of education for analyses involving processing speed. These covariates are frequently employed in studies of CSVD due to their relevance to cognitive impairment (Olivia K. L. Hamilton *et al*., 2021). The statistical significance level was set at α < 0.05, with correction for multiple comparisons using the false discovery rate (FDR) method (the Benjamini and Hochberg procedure) (Benjamini and Hochberg, 1995).

#### 2.5.1 Effect of WMH burden on white matter microstructure

First, we investigated the microstructure of the SWM (and DWM) under increasing WMH burden. WMH-total volume was set as the independent variable, and we investigated its associations with the four diffusion measures in the SWM and DWM using simple linear regression analyses. There were 8 comparisons in this experiment, and results were corrected for multiple comparisons using FDR. The standardized β coefficient of each variable was reported for each regression model.

#### 2.5.2 Association of each imaging marker with processing speed

Second, we investigated if imaging markers in the SWM (and DWM) were associated with processing speed in CSVD. Imaging markers included WMH-total, WMH-SWM, and WMH-DWM volumes, as well as the four diffusion measures in the SWM and DWM. To do so, we first assessed the association between each imaging marker and processing speed using linear regression analyses. In each regression model, the imaging marker was set as the independent variable, and the TMT-A completion time was set as the dependent variable. All results were corrected for the number of comparisons (3 comparisons for the WMH volume analyses, and 8 comparisons for the diffusion measure analyses) using the FDR method.

In addition to the association analysis for each image marker independently, we also performed a multivariable regression analysis to assess the explanatory power of all imaging markers together. Considering the presence of potential collinearity among these imaging markers (see Supplementary Figure S1 for intercorrelations between all variables using Pearson correlation analysis), we performed random forest regression with conditional inference trees (Strobl *et al*., 2007), an advanced machine learning based technique to estimate importance of each variable while accounting for all other variables. Compared to the commonly used multiple linear regression, conditional forest regression offers increased robustness against multicollinearity (Hothorn, Hornik and Zeileis, 2006) and is unbiased when subsampling, avoiding potentially biased estimates in the presence of high interactions among different types of predictions (such as categorical and continuous variables) (Strobl *et al*., 2007). Conditional forest regression has been successfully applied in dMRI studies to assess the contributions of multiple imaging markers (Pechtel *et al*., 2014; Finsterwalder *et al*., 2020; Nemy *et al*., 2020; Konieczny *et al*., 2021), including dealing with cognition in association with multiple microstructural measures (Konieczny *et al*., 2021) and diffusion measures of different tracts (Nemy *et al*., 2020), as well as associations among multiple imaging markers (Finsterwalder *et al*., 2020). The R package ‘party’ (version 1.3–7) was applied (Strobl *et al*., 2007). We calculated 1501 conditional inference trees with unbiased variable selection (unbiased resampling scheme) (Hothorn, Hornik and Zeileis, 2006) using standard parameters (5 randomly preselected variables for each split). From these trees, we next calculated a conditional permutation importance from 400 repetitions (Duering *et al*., 2018). The resulting importance of each variable was calculated as the increase in the mean squared error (MSE) of the predicted response variable (i.e., TMT-A completion time in our study), where an important variable would produce a large increase in MSE (Strobl *et al*., 2007). To further investigate the robustness of the results from the conditional forest regression, we also performed a standard random forest regression that did not include conditional inference trees. Results are reported in Supplementary Methods, where we showed that the overall importance ranking of variables corresponded across both analyses (Figure S2 in Supplementary and Figure 2 in Results).

**Figure 2.**
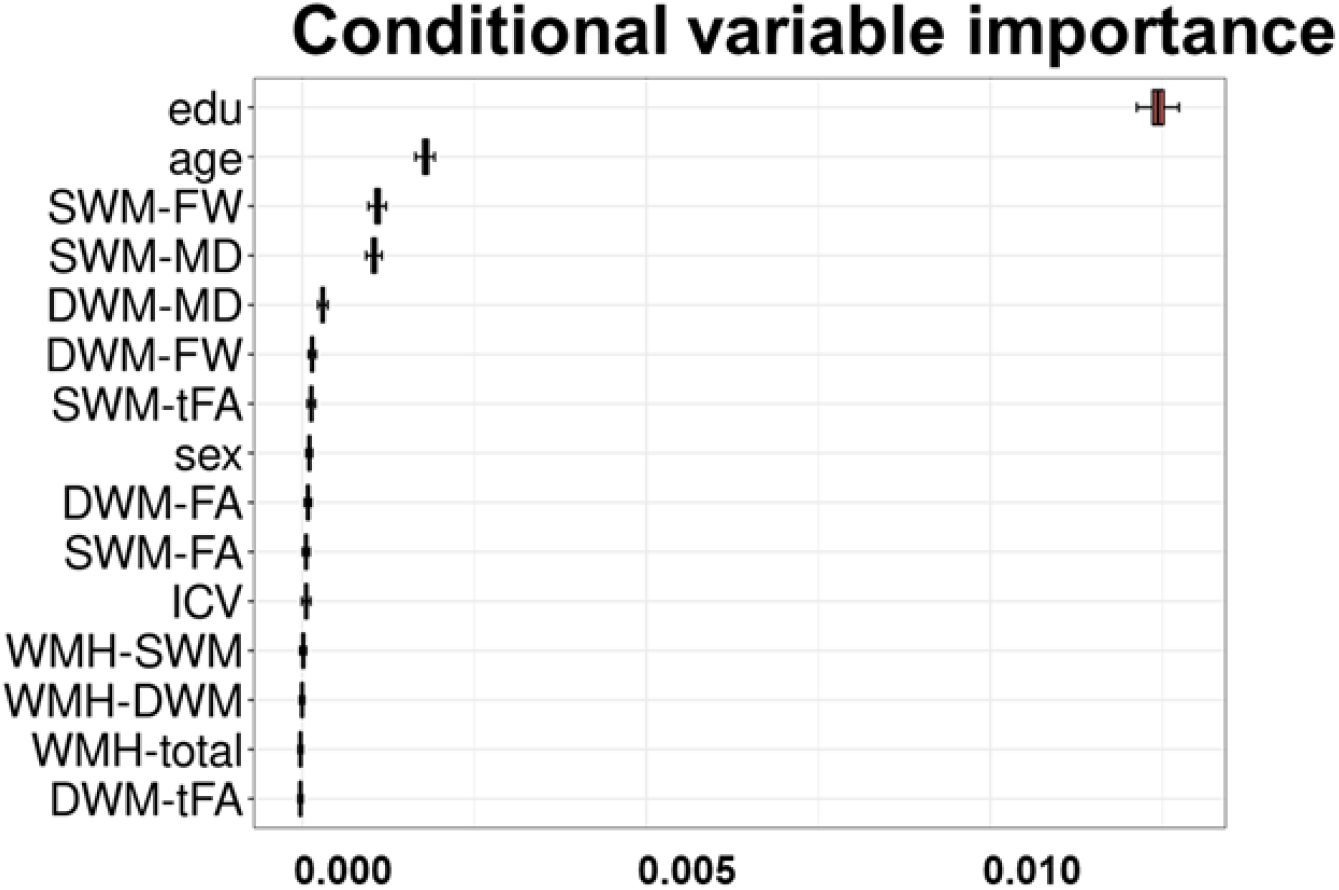
Conditional importance of each variable with regard to processing speed (dependent variable) computed using the conditional random forest regression. Plot shows the conditional variable importance with mean values and 95% confidence intervals calculated from 400 repetitions. Variable importance is calculated as the increase in the MSE of the predicted response variable, where a larger value indicates a higher variable importance. *Abbreviations*: MSE, mean squared error; SWM, superficial white matter; DWM, deep white matter; WMH, white matter hyperintensity; FA, fractional anisotropy; MD, mean diffusivity; FW, free-water; FAt, FW-corrected FA.

To assess the effect of regional SWM microstructure on processing speed, we examined associations between processing speed and the four diffusion measures in each of 8 lobar SWM regions using simple linear regression analyses. The standardized β coefficient of each variable was estimated for each regression model. Results were corrected for 32 (4*8) comparisons using the FDR method.

#### 2.5.3 Abnormalities of SWM and DWM in different levels of disease

Third, we assessed microstructures in the SWM (and DWM) and cognitive performance in different levels of disease. The subject sample was split into 4 subgroups with different levels of WMH burden, according to the quartiles of WMH-total volume (the first quartile and the fourth quartile of patients had the lowest and the highest WMH burden, respectively). We compared demographic and clinical characteristics among the four subgroups using the chi-square test, one-way analysis of variance (ANOVA) or Kruskal-Wallis test, depending on the type and distribution of dependent variables. Imaging markers in the SWM and the DWM were calculated in each subgroup. The processing speed measures and diffusion measures were compared among the four subgroups using ANOVA and Tukey’s post hoc tests. We identified the disease levels in which processing speed and imaging markers had significant between-group differences.

#### 2.5.4 Mediation effect of SWM between WMH burden and processing speed

Fourth, we tested whether the relationship between WMH volume and processing speed was mediated by regional FW, i.e., the diffusion measure with the highest association with WMH volume (according to the analysis in 2.5.1) and the highest association with processing speed (according to the analysis in 2.5.2). To do so, we performed mediation analysis (Liu *et al*., 2016; Avram *et al*., 2019; Brown *et al*., 2019) using PROCESS version 4.0 (https://processmacro.org/index.html) in SPSS (Hayes, 2017). FW of SWM and DWM were introduced as two mediators between WMH volume and processing speed. We examined whether the total effect (c) of WMH volume on processing speed is due to a significant direct effect (c’) or instead is explained by indirect effects (ab) through mediator variables. Effect sizes were calculated as standardized regression coefficients (Hayes, 2017; Su *et al*., 2018; Brown *et al*., 2019). A bootstrapping approach with 5000 iterations was used to estimate confidence intervals (CI) (Gazes *et al*., 2016; Brown *et al*., 2019). Effects with bootstrapped 95% CI not including zero were considered significant (Gazes *et al*., 2016; Avram *et al*., 2019; Brown *et al*., 2019).

## 3. Results

A total of 141 subjects with CSVD were included in the final analysis. Demographic, clinical and MRI characteristics of subjects are provided in Table 1.

**Table 1.**
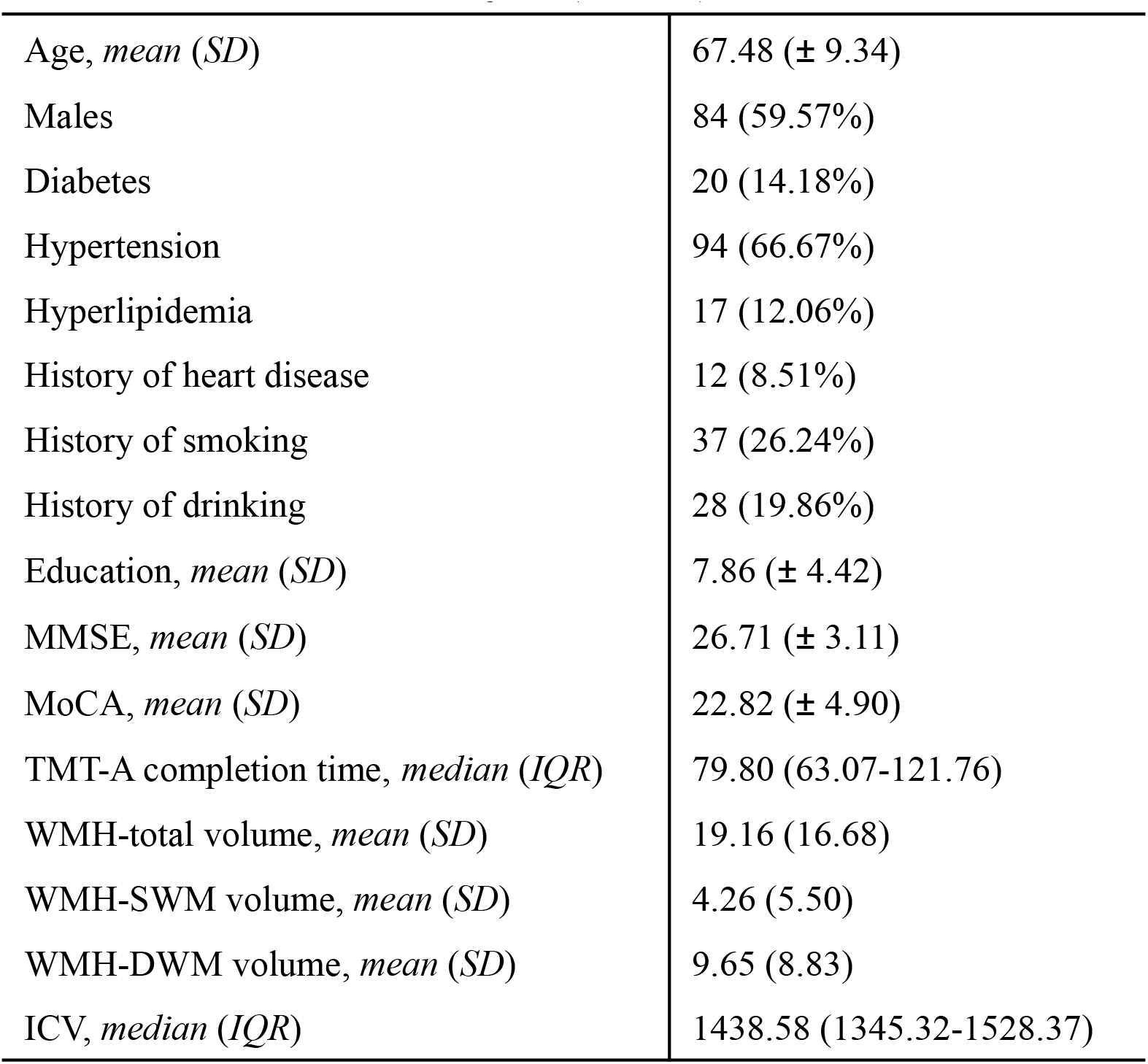
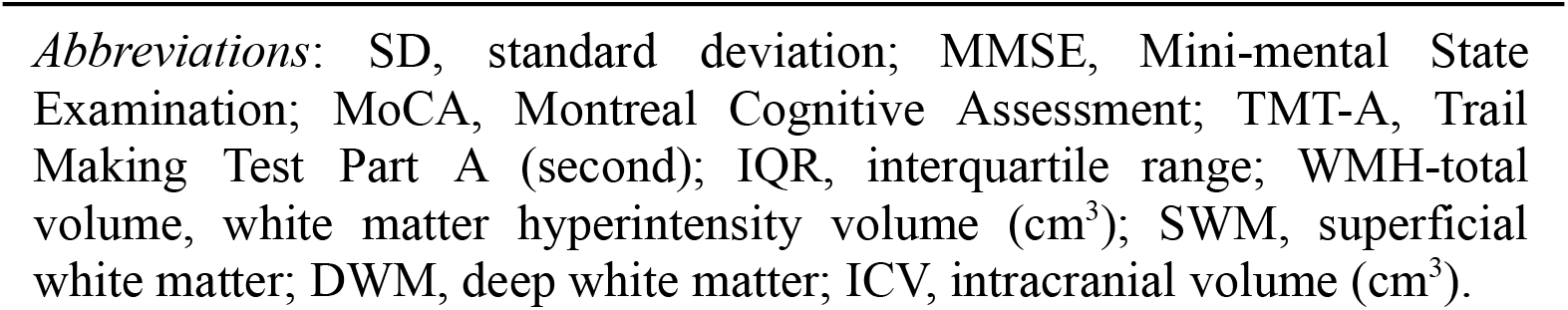
Characteristics of subjects (N = 141)

|  |  |
| --- | --- |
| Age, *mean (SD)* | 67.48 (± 9.34) |
| Males | 84 (59.57%) |
| Diabetes | 20 (14.18%) |
| Hypertension | 94 (66.67%) |
| Hyperlipidemia | 17 (12.06%) |
| History of heart disease | 12 (8.51%) |
| History of smoking | 37 (26.24%) |
| History of drinking | 28 (19.86%) |
| Education, *mean (SD)* | 7.86 (± 4.42) |
| MMSE, *mean (SD)* | 26.71 (± 3.11) |
| MoCA, *mean (SD)* | 22.82 (± 4.90) |
| TMT-A completion time, *median (IQR)* | 79.80 (63.07-121.76) |
| WMH-total volume, *mean (SD)* | 19.16 (16.68) |
| WMH-SWM volume, *mean (SD)* | 4.26 (5.50) |
| WMH-DWM volume, *mean (SD)* | 9.65 (8.83) |
| ICV, *median (IQR)* | 1438.58 (1345.32-1528.37) |

*Abbreviations:* SD, standard deviation; MMSE, Mini-mental State Examination; MoCA, Montreal Cognitive Assessment; TMT-A, Trail Making Test Part A (second); IQR, interquartile range; WMH-total volume, white matter hyperintensity volume (cm<sup>3</sup>); SWM, superficial white matter; DWM, deep white matter; ICV, intracranial volume (cm<sup>3</sup>).

### 3.1 FW has the strongest association with WMH burden

To characterize the microstructure in the SWM (and in the DWM) under increasing WM burden, associations between regional diffusion measures and WMH volume were analyzed. FA, MD and FW in SWM and DWM were significantly associated with WMH-total volume (all *p*_*FDR*_ <0.001, Table 2). As the overall disease burden increased (as measured by the WMH-total volume), FA decreased, while MD and FW increased. The FW measure had the strongest association with WMH-total volume in both SWM and DWM. The disease burden (as measured by WMH-total volume) had the largest effect on diffusion measures in the DWM (where a larger lesion volume was observed; see WMH-DWM in Table 1). This effect is indicated by the larger magnitude of the standardized β coefficients of the diffusion measures in the DWM, in comparison to those of the SWM. The free-water-corrected diffusion measure FAt showed no significant association with the WMH-total volume.

**Table 2.** Associations between regional diffusion measures and WMH-total volume.

| Diffusion measures | SWM |  | DWM |  |
| --- | --- | --- | --- | --- |
| | $\beta_s$ | $p$ | $\beta_s$ | $p$ |
| **FW** | 0.518 | **<0.001** | 0.838 | **<0.001** |
| **MD** | 0.459 | **<0.001** | 0.752 | **<0.001** |
| **FA<sub>t</sub>** | 0.020 | 0.791 | -0.083 | 0.368 |
| **FA** | -0.362 | **<0.001** | -0.664 | **<0.001** |

Note: Adjusted for age, sex and ICV. Significant FDR-corrected results are in BOLD.
*Abbreviations:* $\beta_s$ , standardized $\beta$ coefficient; WMH, white matter hyperintensity; SWM, superficial white matter; DWM, deep white matter; FA, fractional anisotropy; MD, mean diffusivity; FW, free-water; FA<sub>t</sub>, FW-corrected FA; ICV, intracranial volume.

### 3.2 Association of each imaging marker with processing speed

Next, we investigated if imaging markers in the SWM (and DWM) were associated with processing speed in CSVD.

#### 3.2.1 Imaging markers in SWM and DWM are associated with processing speed

As shown in Table 3, the WMH burden measures were all significantly associated with processing speed, with similar standardized β coefficients for all regressions.

**Table 3.** Associations between global and regional WMH volumes and processing speed.

| WMH volume | $\beta_s$ | $p$ |
| --- | --- | --- |
| WMH-total | 0.154 | **0.020** |
| WMH-SWM | 0.153 | **0.020** |
| WMH-DWM | 0.146 | **0.020** |

Note: The association analyses are adjusted for age, sex, ICV and education. Significant FDR-corrected results are in BOLD.
*Abbreviations:* $\beta_s$ , standardized $\beta$ coefficient; SWM, superficial white matter; DWM, deep white matter; WMH, white matter hyperintensity; ICV, intracranial volume.

Furthermore, SWM and DWM diffusion measures (FA, MD and FW) were significantly associated with processing speed (Table 4). As expected, the poorer the cognitive performance, the lower the FA and the higher the MD and FW. SWM FW was most strongly associated with processing speed, with the largest magnitude standardized β coefficient (β_S_ = 0.289), followed by SWM MD (β_S_ = 0.246). No significant association was found between free-water-corrected diffusion measure FAt and processing speed in SWM or DWM.

**Table 4.** Associations between each diffusion measure and processing speed Diffusion measures.

| Diffusion measures | SWM |  | DWM |  |
| --- | --- | --- | --- | --- |
| | $\beta_s$ | $p$ | $\beta_s$ | $p$ |
| FW | 0.289 | **<0.001** | 0.240 | **<0.001** |
| MD | 0.246 | **<0.001** | 0.245 | **<0.001** |
| FAt | -0.003 | 0.962 | -0.027 | 0.771 |
| FA | -0.179 | **0.006** | -0.186 | **0.004** |

Note: Adjusted for age, sex, ICV and education. Significant FDR-corrected results are in BOLD.
*Abbreviations:* $\beta_s$ , standardized $\beta$ coefficient; DWM, deep white matter; SWM, superficial white matter; WMH, white matter hyperintensity; FA, fractional anisotropy; MD, mean diffusivity; FW, free-water; FAt, FW-corrected FA; ICV, intracranial volume.

#### 3.2.2 FW in the SWM contributes most to processing speed in multivariable analysis

To assess the contribution of each imaging marker to processing speed while accounting for multicollinearity (see Supplementary Figure S1 for a matrix of correlations between all variables), we applied conditional random forest regression and calculated the conditional variable importance. Across all imaging markers, significant variables included FW and MD in the SWM, showing high contributions to processing speed (Fig. 2). Specifically, the FW in the SWM had the highest variable importance.

#### 3.2.3 Diffusion measures of SWM in different brain lobes are widely associated with processing speed

To investigate any regional effects of SWM microstructure on processing speed, we examined associations between processing speed and lobar SWM microstructure. In all lobar SWM regions, FW and MD were strongly associated (p < 0.001) with processing speed (Supplementary Table S2). In contrast, associations between lobar SWM FA and processing speed were found only in the left frontal and right parietal SWM. No significant association was found for FAt.

### 3.3 Comparison of imaging markers of SWM and DWM between different levels of disease

The study sample was split into quartiles according to the total WMH volume, producing 4 subgroups with different levels of WMH burden. There was no significant difference in age, sex, ICV or education between subgroups (See Supplementary Table S1 for this information for each group).

Figure 3(a) shows the TMT-A completion time and regional WMH volumes for subgroups of different WMH burden levels. The WMH-DWM burden significantly increased at all levels of disease burden (i.e. between all compared groups: Groups 1 and 2, Groups 2 and 3, and Groups 3 and 4). However, a significant increase in WMH-SWM was found only between higher levels of disease burden (between Groups 3 and 4). This was accompanied by a significant increase in TMT-A completion time (a decrease in processing speed) at higher levels of disease burden (between Groups 3 and 4).

**Figure 3.**
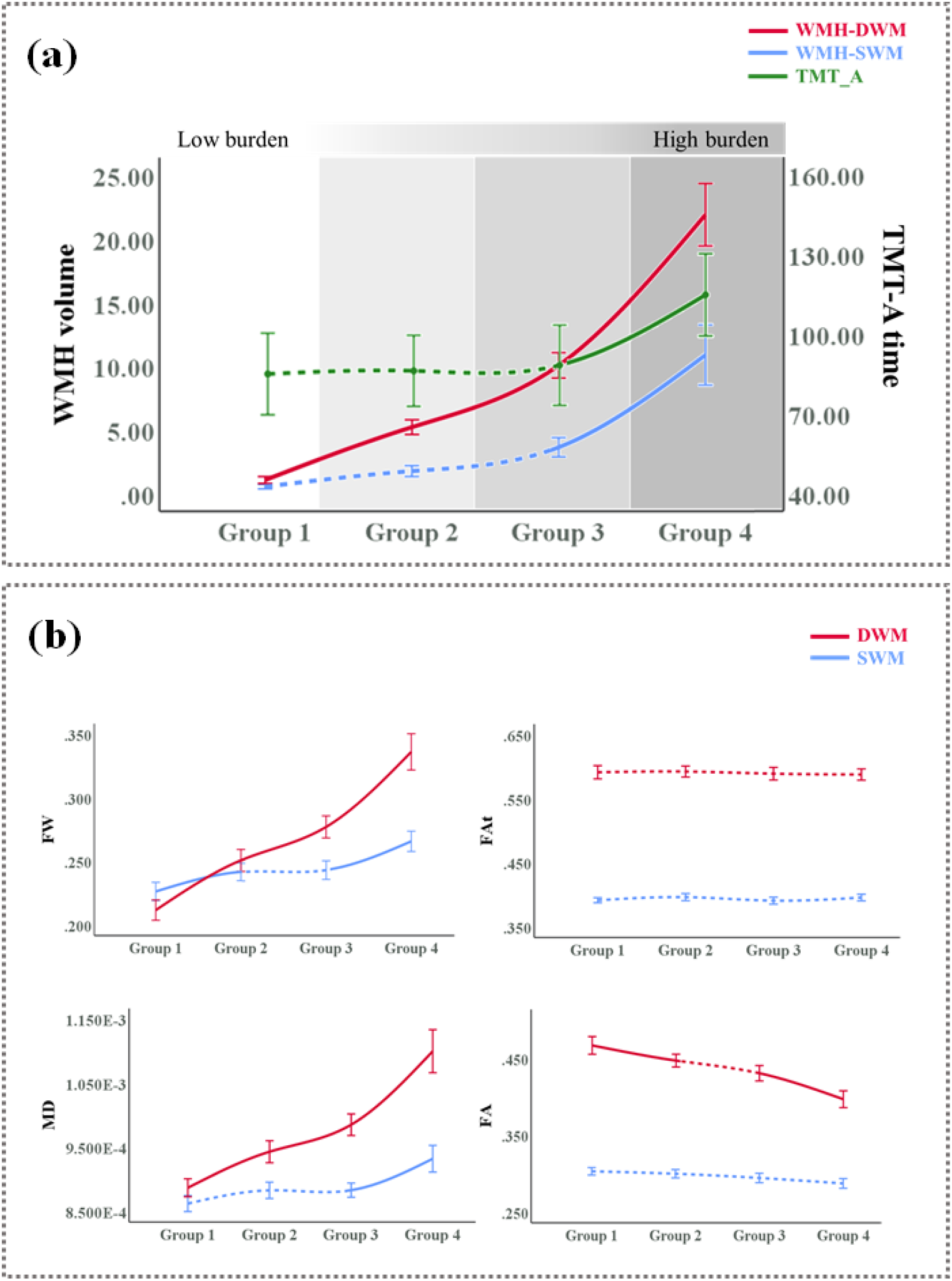
The TMT-A completion time and imaging markers in subgroups of different WMH burden levels. (a) The TMT-A completion time and regional WMH volumes. The left y-axis represents the WMH volume, and the right y-axis represents the completion time of TMT-A. (b) Diffusion measures in the SWM and the DWM. Patients in Group 1 and 4 had the lowest and highest level of WMH burden, respectively. Between-group differences were compared. Solid line segments indicate that the diffusion measures significantly differed compared to the previous group, while dashed line segments indicate that between-group differences did not reach a statistically significant level. *Abbreviations*: DWM, deep white matter; SWM, superficial white matter; WMH, white matter hyperintensity; FW, free-water; FA, fractional anisotropy; MD, mean diffusivity; FAt, FW-corrected FA.

**Figure 4.**
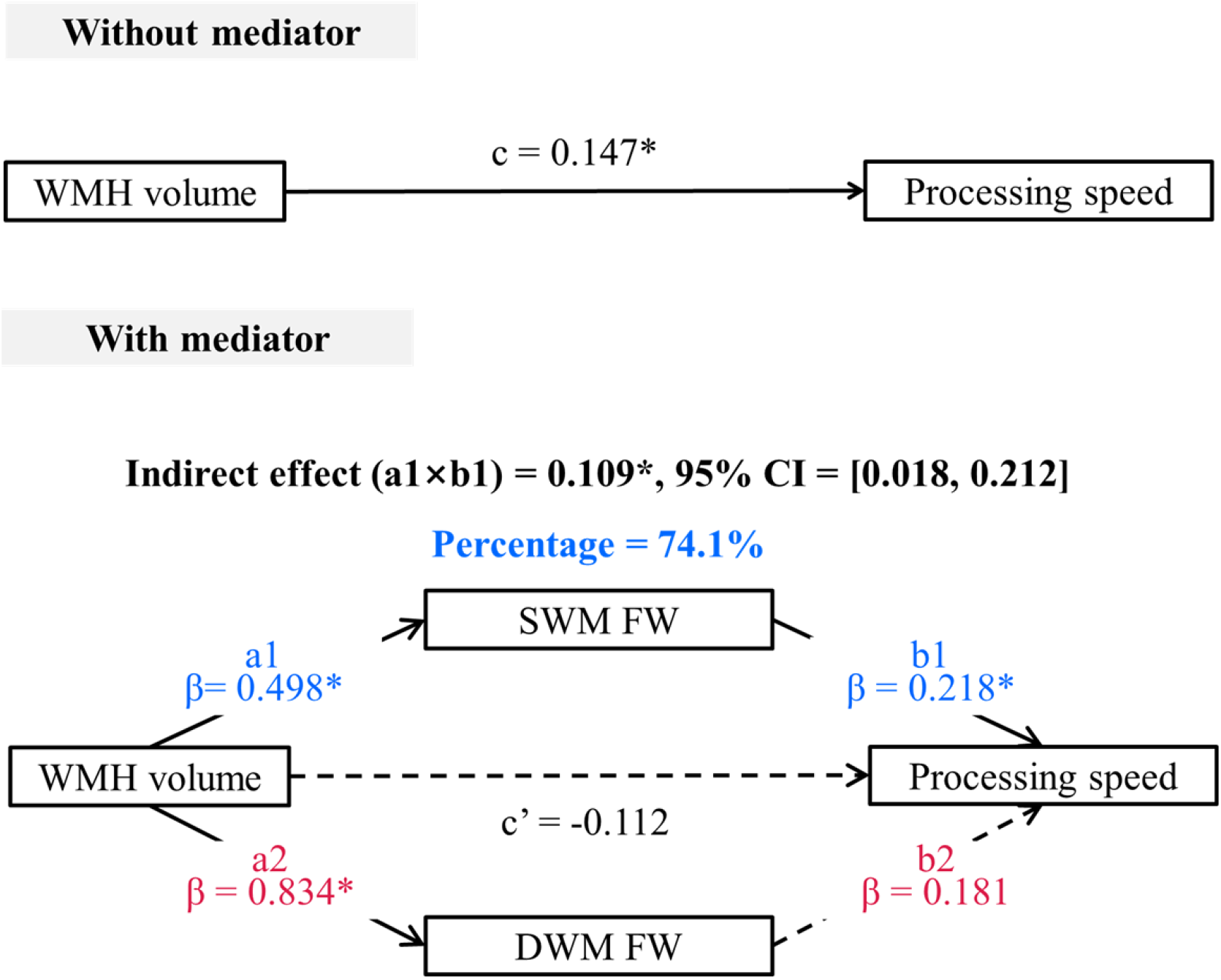
Mediation of regional FW between WMH volume and processing speed. The effect of WMH volume on processing speed is shown by the direct effect (c) without and the indirect effect (c’) with mediators. Standardized β-coefficients of each path (a and b) are shown for two mediators (*p < 0.05). Significant paths are indicated by solid arrows, and non-significant paths are shown by dashed arrows. Indirect effects are statistically significant at the 95% CI when the CI does not include 0. As the direct effect (c’) is not significant, SWM FW fully mediates the relationship between WMH volume and processing speed. (This analysis controlled for age, sex, ICV and education.) *Abbreviations*: WMH, white matter hyperintensity; SWM, superficial white matter; DWM, deep white matter; FW, free-water; CI, confidence interval.

Figure 3(b) shows all diffusion measures in the SWM and DWM in the subgroups of different WMH burden levels. DWM-FW and DWM-MD significantly increased at all levels of disease burden (i.e. between all compared groups: Groups 1 and 2, Groups 2 and 3, and Groups 3 and 4). The overall trend of DWM-FA was downwards, though not all decreases between groups were significant. In contrast, the between-group differences in diffusion measures in the SWM were less pronounced than those in the DWM. Specifically, only SWM-FW significantly differed in subjects with relatively lower WMH burdens (between Groups 1 and 2), suggesting that the SWM-FW measure is more sensitive than SWM-MD or SWM-FA. Significant increases in SWM-FW and SWM-MD could be observed at higher levels of disease burden (between Groups 3 and 4). Between-group differences in FAt were relatively slight at all levels of disease burden and not significant in the SWM or the DWM.

### 3.4 Free water in the SWM fully mediates the WMH volume effect on processing speed

Given significant effects of WMH volume and regional FW on processing speed, we performed mediation analysis to further determine whether FW in the SWM and the DWM could mediate the association between WMH volume and processing speed. Mediation analysis revealed a significant indirect effect of WMH volume on processing speed via SWM FW (a1×b1 = 0.109, the bootstrapped 95% CI = [0.018, 0.212]). As a mediator, SWM FW could account for 74.1% of the total effect (c). For FW in the DWM, the bootstrapped 95% CI included zero ([-0.124, 0.450]), indicating that the mediated effect was not significant. The direct effect of WMH volume on processing speed was not significant after adjusting for the mediators (c’ = − 0.112, p > 0.05), supporting a full mediation model. These results suggest that SWM FW fully mediates the relationship between the WMH volume and processing speed.

## 4. Discussion

This study applies multiple imaging markers to assess white matter abnormalities in the SWM and the DWM, and investigates their contributions to processing speed in CSVD. The main findings can be summarized as follows: i) the contributions to processing speed of the SWM (as reflected in imaging markers) are comparable to those of the DWM, despite the fact that the SWM has a lower WMH burden than the DWM; ii) SWM FW has the strongest association with processing speed among all imaging markers and, unlike the other diffusion MRI measures, significantly increases in subjects with low WMH burden (possibly representing early stages of disease); iii) under high burden of disease, the involvement of WMH in the SWM is accompanied by significant differences in processing speed and white matter microstructure; iv) the effect of WMH volume on processing speed is fully mediated by SWM FW, and no mediation effect of DWM FW is observed. Our findings provide new insights into processing speed in patients with CSVD and suggest the SWM as a potential target for future research.

Our results extend the present literature on white matter abnormalities and their impacts on processing speed in CSVD, providing evidence for the importance of the SWM. A number of CSVD studies have found strong associations of cognitive performance with diffusion MRI measures in the DWM (O’Sullivan *et al*., 2005; Quinque *et al*., 2012; Tuladhar *et al*., 2015; Chen *et al*., 2019; Huang *et al*., 2020; Liu *et al*., 2020; Vergoossen *et al*., 2021) and in white matter areas without visible WMH (M. O’Sullivan *et al*., 2004; M. O’Sullivan *et al*., 2004; Schmidt *et al*., 2010; Jokinen *et al*., 2013; Lawrence *et al*., 2013, 2014; Tuladhar *et al*., 2015; Zhang, Wong, Uiterwijk, *et al*., 2017; Tozer *et al*., 2018). Although postmortem histological studies have observed white matter abnormalities in the SWM in CSVD (Humphreys, Smith and Wardlaw, 2021), limited dMRI studies have investigated the role of the SWM in CSVD, showing a reduction in SWM connectivity (Lawrence *et al*., 2014; Tuladhar *et al*., 2017; van Leijsen *et al*., 2019). The present study suggests that processing speed in CSVD is particularly related to microstructural abnormalities in the SWM. The significance of the SWM for cognitive performance, especially in executive function, processing speed and visuomotor-attention, has been reported in aging (Nazeri *et al*., 2015), Alzheimer’s disease (Phillips *et al*., 2016; Reginold *et al*., 2016) and psychiatric disorders (Nazeri *et al*., 2013; D’Albis *et al*., 2017). In the current study, we mainly focused on the processing speed, which is the primary cognitive dysfunction in CSVD (Peng, Geriatric Neurology Group, Chinese Society of Geriatrics and Clinical Practice Guideline for Cognitive Impairment of Cerebral Small Vessel Disease Writing Group, 2019; Konieczny *et al*., 2021). We found that processing speed was significantly associated with SWM abnormalities measured by imaging markers, including SWM FA, MD, FW and WMH-SWM. Interestingly, the contributions of SWM imaging markers to processing speed were higher than those of the DWM in multivariable analyses (Result 3.3), despite the fact that the SWM had a lower WMH burden, and diffusion measures in the SWM were less affected by the total WMH burden (see the magnitude of the standardized β coefficients, Table 2). This finding indicates that SWM may play a role of limited change but significant effect in cognition. In our study, the potential importance of SWM in CSVD is further emphasized by the result that SWM FW fully mediates the effect of WMH on processing speed. In related work, mediation analyses have shown that global dMRI measures (including FW measured in the entire white matter skeleton (Maillard *et al*., 2019) and local network efficiency, a property of the entire white matter connectome (Vergoossen *et al*., 2021)) mediate the relationship between WMH volume and cognitive performance measures in aging. Overall, our results demonstrate the importance of studying localized dMRI measures in the SWM and suggest that the importance of the SWM may be underestimated in previous studies of CSVD.

SWM imaging markers such as those studied here, in conjunction with previously studied DWM imaging markers, may provide useful information about the disease course in CSVD. Previous CSVD research has linked disease severity to abnormalities in regional white matter (e.g., DWM, normal-appearing white matter and peripheral white matter connections) that were associated with the severity of persisting cognitive impairments (Jokinen *et al*., 2013; Schmidt, Seiler and Loitfelder, 2016; van Leijsen *et al*., 2017, 2019). In the present study, we identified the WMH burden levels in which processing speed and imaging markers had significant between-group differences. It is worth noting that while the WMH-DWM burden increased from Group 1 to 4, only between Groups 3 and 4 (with higher WMH burden) did we find significant WMH-SWM and processing speed differences (Fig. 3). Our findings suggest that cognitive performance may be related to the involvement of WMH-SWM. This result further shows that the contribution of WMH-SWM to cognitive performance may be a late event. In addition, close associations between microstructural diffusion measures of DWM and SWM (Figure S1) suggest that earlier microstructural abnormalities in the DWM might predict the appearance of later changes in the SWM, which may be an important cumulative factor for a steep difference in processing speed between two groups in the later stages. Considering that the SWM has the above-mentioned characteristics of limited changes but significant effect on cognitive performance, the detection of subtle changes in the SWM is very important. Our results show that SWM microstructural differences can be detected by FW in the scenario of low WMH burden, which could not be achieved by conventional diffusion measures (Fig. 3). Therefore, it is possible that SWM imaging markers may be useful for monitoring the disease course, from early SWM microstructural change (potentially characterized by SWM FW) to late pronounced difference in cognitive performance (potentially characterized by WMH-SWM).

The present study suggests that FW in the SWM may constitute an early marker of white matter microstructural abnormality in CSVD. Although FW was originally estimated to correct for freely diffusing extracellular water (Pasternak *et al*., 2009), many studies have found that FW is particularly sensitive to disease severity (Ji *et al*., 2017; Duering *et al*., 2018; Maillard *et al*., 2019; Gullett *et al*., 2020; Huang, Zhang, Jiaerken, Wang, Hong, *et al*., 2021; Yu *et al*., 2021). Using FW imaging, increasing evidence suggests a prominent increase in extracellular water in CSVD, with less demyelination and axonal loss than would be suggested by conventional diffusion measures (Duering *et al*., 2018; Wardlaw, Smith and Dichgans, 2019). In line with these previous findings, we found that extracellular water in the DWM (DWM-FW) significantly increased in the subgroups with low WMH burden. In addition, our results extended this finding to the SWM, showing that, among all the imaging markers studied, FW was the most sensitive to SWM microstructure. Of note, a significant increase in SWM-FW could be observed in the subgroups with low WMH burden (Groups 1 and 2; Figure 3), while WMH-SWM only slightly increased between these two groups. This finding is consistent with the current view, which suggests that the increased water content in white matter is associated with the early stage of WMH formation (Muñoz Maniega *et al*., 2017; Wardlaw, Smith and Dichgans, 2019; Huang, Zhang, Jiaerken, Wang, Yu, *et al*., 2021). Our results indicate that increased extracellular water, as measured by FW, may be an early sign and main process for SWM microstructural abnormality under low WMH burden.

To our knowledge, FW in the SWM in CSVD was not investigated before, and therefore the factors driving increased SWM FW remain to be determined. Although the exact pathology contributing to the diffusion changes cannot be determined in the current study, there are several potential pathological processes relevant for CSVD that could explain the increased extracellular FW in the SWM. One possible factor is myelin vacuolization, which contributes to white matter rarefaction, and vacuoles in white matter enable interstitial fluid to accumulate (Murray *et al*., 2012), further increasing the extracellular water. In an animal study of genetic CSVD (Cognat *et al*., 2014), segmental vacuolization of the white matter was reported as the earliest pathological change. In humans, a postmortem study from Erkinjuntti et al., has found vacuolization in the SWM in nearly half of patients with CSVD-related vascular disorders (Erkinjuntti *et al*., 1996). This evidence suggests that vacuolization in the SWM may be a pathological process related to small vessel disease. An alternative pathological process that may drive FW is the edema caused by BBB breakdown in WMH, which can increase extracellular water (Duering *et al*., 2018; Di Biase, Chad and Pasternak, 2021). It needs to be noted that the aforementioned myelin vacuolization can promote BBB leakage (Cognat *et al*., 2014). Therefore, the involvement of BBB damage in the SWM may be in the later stages of the disease, which may be a contributor to a steep increase in FW observed in this study (Fig. 3, Groups 3 and 4). Another mechanism receiving recent attention is glymphatic dysfunction in CSVD (Benveniste and Nedergaard, 2021). Recent studies found that the impaired drainage of interstitial fluid can increase FW around the perivascular space (Huang, Zhang, Jiaerken, Wang, Yu, *et al*., 2021; Jiaerken *et al*., 2021), which is the fluid-filled space surrounding blood vessels (Wardlaw *et al*., 2020). The increased water content in the dilated perivascular space may be related to WMH formation (Weller *et al*., 2015; Huang, Zhang, Jiaerken, Wang, Yu, *et al*., 2021). Although this question did not constitute the objective of our study, it should be noted that the increased water content in the dilated perivascular spaces may partly contribute to the observed increased FW in the SWM. Taken together, our study suggests that SWM FW may be a valuable marker for CSVD.

Our study shifts the focus of CSVD-related white matter abnormalities to the SWM region, and the results highlight the role of the SWM in CSVD-related dysfunction. Some limitations of the study are as follows. First, although the present study provides evidence supporting the role of the SWM in CSVD, the cross-sectional design limits any exploration of longitudinal SWM change and its effects on cognitive decline. Future longitudinal studies of CSVD are needed. Second, although processing speed is the primary cognitive function affected in CSVD, other neuropsychological domains, such as memory and depression, need to be investigated in the future. Third, while we have investigated the effects of CSVD on the SWM microstructure as a whole, future work may shed light on the contribution of localized WMH to these changes, and any effect of CSVD on normal-appearing white matter in the SWM. Fourth, cortical atrophy may contribute to cognitive performance, and its association with abnormalities in the SWM needs to be explored in the future. Finally, future studies may apply more advanced diffusion acquisitions such as multi-shell to improve robustness of the model fit, and also apply tractography to identify the white matter tracts in the SWM, which may provide a more precise assessment for microstructural changes in the SWM (Zhang *et al*., 2018; Guevara *et al*., 2020).

## 5. Conclusion

In this study, our findings identify SWM abnormalities in CSVD and suggest that the SWM has an important contribution to processing speed. Our results suggest that processing speed decreases in CSVD may be driven by the involvement of WMH lesions in the SWM. Results indicate that SWM FW fully modulates the association between WMH burden and processing speed, while no mediation effect of DWM FW was observed. Overall, FW in the SWM is a sensitive marker of microstructural changes associated with cognition in CSVD. This study extends the current understanding of CSVD-related dysfunction and suggests that the SWM, as an understudied region, can be a potential target for monitoring pathophysiological processes in future research.

## 6. Acknowledgements

The authors gratefully acknowledge the following funding grants:

RK: P41 EB015902 (NAC), P41EB028741 (AT-NCIGT), R01 CA235589 (LNQ), National Cancer Data Ecosystem, Task Order No. 413 HHSN26110071 under Contract No. HHSN261201500003l; LJO and FZ: R01MH119222, R01MH125860, P41EB015902, R01MH074794; FZ also acknowledges a BWH Radiology Research Pilot Grant Award; NM: R01MH125860, R01MH112748, R01MH111917, K24MH116366, R01AG042512, R21DA042271; MZ: the National Natural Science Foundation of China (Grant Nos. 81971577, 81771820, 82101987, 81901706 & 82101984), and the Natural Science Foundation of Zhejiang Province (Grant No. LSZ19H180001), the China Postdoctoral Science Foundation (Grant No. 2019M662083) and the Zhejiang province Postdoctoral Science Foundation.

## Author contribution statement

**Shuyue Wang:** Conceptualization, Methodology, Writing - review & editing, Data curation, Data acquisition. **Fan Zhang:** Conceptualization, Methodology, Writing - review & editing, Data curation, Software. **Peiyu Huang:** Conceptualization, Writing - review & editing. **Hui Hong:** Conceptualization, Writing - review & editing, Data acquisition. **Yeerfan Jiaerken:** Conceptualization, Writing - review & editing, Data acquisition. **Xinfeng Yu:** Conceptualization, Writing - review & editing, Data acquisition. **Ruiting Zhang:** Conceptualization, Writing - review & editing, Data acquisition. **Qingze Zeng:** Conceptualization, Writing - review & editing. **Yao Zhang:** Conceptualization, Writing - review & editing, Data acquisition. **Ron Kikinis:** Conceptualization, Writing - review & editing. **Yogesh Rathi:** Conceptualization, Writing - review & editing. **Nikos Makris:** Conceptualization, Writing - review & editing. **Min Lou:** Conceptualization, Writing - review & editing, Data acquisition, Resources. **Ofer Pasternak:** Conceptualization, Methodology, Writing - review & editing, Software. **Minming Zhang:** Conceptualization, Writing - review & editing, Data curation, Resources. **Lauren J. O’Donnell:** Conceptualization, Methodology, Writing - review & editing, Data curation, Software.

