## Supplementary for "Superficial white matter microstructure affects processing speed in cerebral small vessel disease"

### 1. Supplementary Methods

#### 1.1 Study search strategy and selection for Table 1

Studies were retrieved in November 2021 using a PubMed literature search on “((processing speed) OR (executive function))+((cerebral small vessel disease))+((diffusion magnetic resonance) OR (diffusion MRI) OR (dMRI) OR (diffusion tensor imaging)).”

#### 1.2 Multivariable analysis using a standard random forest regression

To further investigate the robustness of the results from the conditional forest regression, we also performed a standard random forest regression that does not include conditional inference trees. We performed the random forest regression analysis using “rfPermute” (version 2.5; https://cran.r-project.org/web/packages/rfPermute). For each predictor variable, this analysis produced a variable importance, measured as the percentage of increase in mean square error (%IncMSE) of the predicted response variable (i.e., TMT-A completion time), and a p-value to assess the variable’s statistical significance. An important variable produced a large %IncMSE. (See Supplementary Results section 2.4, below, for this result.)

### 2. Supplementary Results

#### 2.1 Characteristics of subgroups

| **Table S1 Characteristics of subgroups** | | | | | |
| --- | --- | --- | --- | --- | --- |
|  | Group 1 | Group 2 | Group 3 | Group 4 | *p* |
| n | 35 | 35 | 35 | 36 | - |
| Age, *mean* (*SD*) | 64.03 (± 7.25) | 68.83 (± 9.30) | 67.23 (±11.17) | 69.78 (±8.58) | 0.051 |
| Males | 16 (45.71%) | 24 (68.57%) | 21 (60.00%) | 23 (63.89%) | 0.240 |
| Diabetes | 6 (17.14%) | 2 (5.71%) | 8 (22.86%) | 4 (11.11%) | 0.223 |
| Hypertension | 18 (51.43%) | 23 (65.71%) | 28 (80.00%) | 25 (69.44%) | 0.052 |
| Hyperlipidemia | 6 (17.14%) | 1 (2.86%) | 5 (14.29%) | 5 (13.89%) | 0.285 |
| History of heart disease | 2 (5.71%) | 2 (5.71%) | 4 (11.43%) | 4 (11.11%) | 0.679 |
| History of smoking | 5 (14.29%) | 9 (25.71%) | 9 (25.71%) | 14 (38.89%) | 0.076 |
| History of drinking | 5 (14.29%) | 9 (25.71%) | 8 (22.86%) | 6 (16.67%) | 0.568 |
| Education, *mean* (*SD*) | 8.17 (± 4.29) | 7.96 (± 4.48) | 8.57 (± 4.33) | 6.78 (± 4.54) | 0.356 |
| MMSE, *mean* (*SD*) | 26.91 (± 3.16) | 27.15 (± 2.62) | 27.24 (± 3.35) | 25.52 (± 3.12) | 0.083 |
| MoCA, *mean* (*SD*) | 23.18 (± 4.68) | 23.65 (± 4.62) | 23.27 (± 5.25) | 21.15 (± 4.87) | 0.154 |
| TMT-A completion time, *median* (*IQR*) | 68.19 (54.42-102.70) | 74.00 (67.00-95.60) | 76.16 (60.10-104.70) | 107.61 (79.43-151.14) | 0.010* c |
| WMH volume, *mean* (*SD*) | 2.86 (± 1.34) | 10.54 (± 2.21) | 19.69 (± 3.54) | 42.86 (± 13.29) | < 0.001* a, b, c |
| ICV, *median* (*IQR*) | 1414.03 (1350.18-1473.58) | 1464.02 (1346.17-1558.40) | 1465.45 (1331.43-1549.54) | 1447.86 (1347.44-1509.36) | 0.451 |
| Note: * ANOVA p < 0.05. Significant post-hoc tests are indicated by letters (a: Group 1 vs. Group 2, b: Group 2 vs. Group 3, c: Group 3 vs. Group 4).  Abbreviations: SD, standard deviation; MMSE, Mini-mental State Examination; MoCA, Montreal Cognitive Assessment; TMT-A, Trail Making Test Part A (second); IQR, interquartile range; WMH volume, white matter hyperintensity volume (cm^3^); ICV, intracranial volume (cm^3^). | | | | | |

#### 2.2 Associations between processing speed and diffusion measures in lobar SWM

| **Table S2 Associations between processing speed and diffusion measures in lobar SWM** | | | | | | | | | |
| --- | --- | --- | --- | --- | --- | --- | --- | --- | --- |
| **Diffusion measures in lobar SWM** | | **FA** | | **MD** | | **FW** | | **FAt** | |
|  |  | **β_S_** | ***p*** | **β_S_** | ***p*** | **β_S_** | ***p*** | **β_S_** | ***p*** |
| **Frontal** | L | -0.145 | **0.039** | 0.257 | **<0.001** | 0.245 | **0.001** | 0.015 | 0.881 |
|  | R | -0.126 | 0.068 | 0.217 | **0.003** | 0.191 | **0.010** | 0.009 | 0.935 |
| **Occipital** | L | -0.113 | 0.117 | 0.231 | **0.003** | 0.239 | **0.002** | 0.014 | 0.881 |
|  | R | -0.148 | **0.045** | 0.275 | **<0.001** | 0.263 | **0.001** | -0.017 | 0.881 |
| **Parietal** | L | -0.126 | 0.082 | 0.233 | **0.001** | 0.272 | **<0.001** | 0.087 | 0.365 |
|  | R | -0.165 | **0.020** | 0.233 | **0.001** | 0.265 | **<0.001** | 0.003 | 0.935 |
| **Temporal** | L | -0.119 | 0.087 | 0.268 | **<0.001** | 0.222 | **0.002** | 0.027 | 0.881 |
|  | R | -0.135 | 0.065 | 0.269 | **<0.001** | 0.226 | **0.002** | 0.018 | 0.881 |
| Note: Adjusted for age, sex, ICV and education. Significant FDR corrected results are in BOLD.  Abbreviations: β_S_, standardized β coefficient; SWM, superficial white matter; FA, fractional anisotropy; MD, mean diffusivity; FW, free-water; FAt, FW-corrected FA; ICV, intracranial volume.  2.3 Correlation between imaging markers | | | | | | | | | |

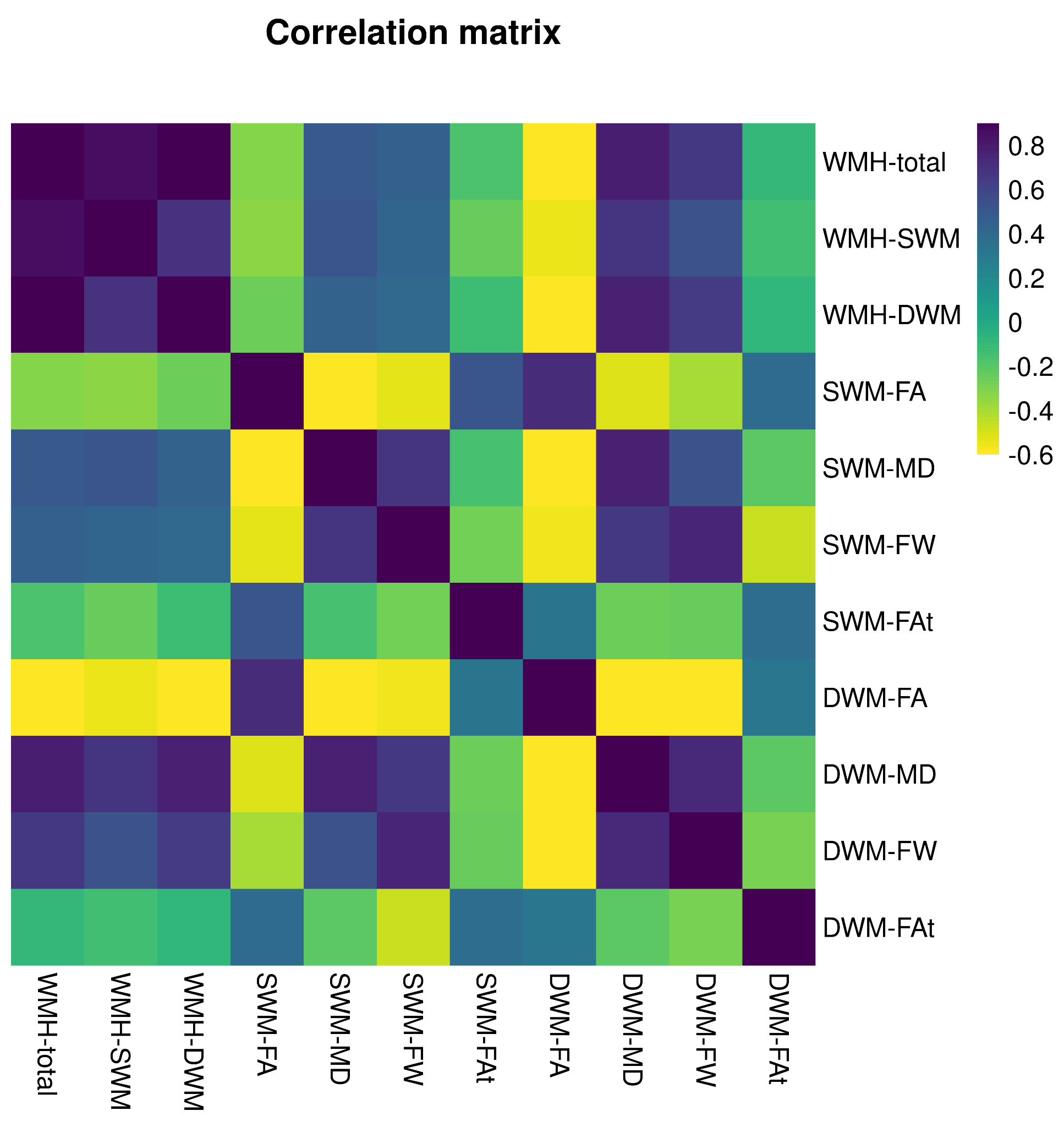

**Figure S1** Correlation matrix illustrating the degree of intercorrelation between imaging markers. Pearson correlation coefficients are coded in color.

*Abbreviations*: DWM, deep white matter; SWM, superficial white matter; WMH, white matter hyperintensity; FA, fractional anisotropy; MD, mean diffusivity; FW, free-water; FAt, FW-corrected FA.

#### 2.4 Result of the multivariable analysis using a standard random forest regression

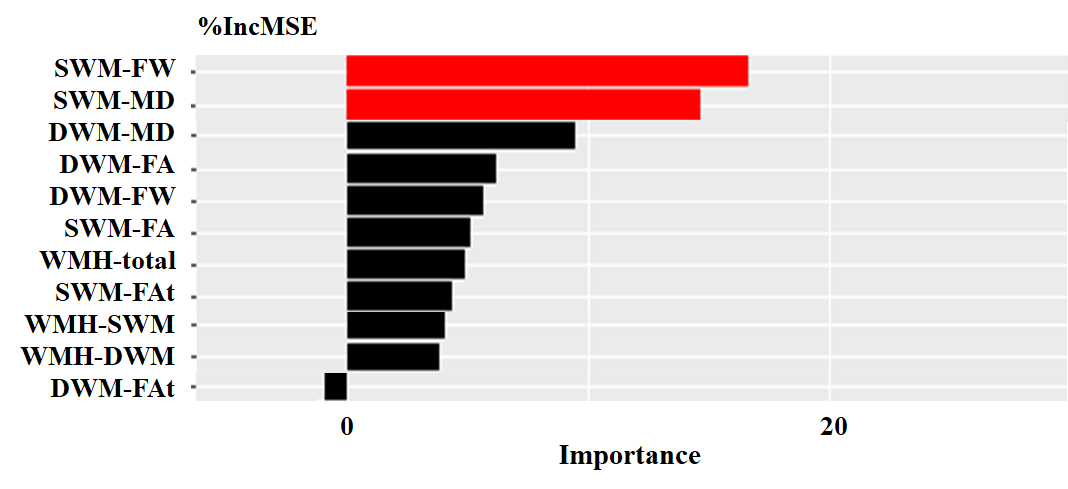

**Figure S2** Multivariable analyses. A standard random forest regression is applied to estimate the importance of independent variables with regard to processing speed. Significant variables are highlighted in red. Variables are sorted by the increase in MSE (%IncMSE). The larger the increase in MSE, the higher importance of the variable.

*Abbreviations*: MSE, mean squared error; SWM, superficial white matter; DWM, deep white matter; WMH, white matter hyperintensity; FA, fractional anisotropy; MD, mean diffusivity; FW, free-water; FAt, FW-corrected FA.
